# ADAM17 is the Principal Ectodomain Sheddase of the EGF-Receptor Ligand Amphiregulin

**DOI:** 10.1101/218891

**Authors:** Vishnu Hosur, Michelle L. Farley, Lisa M. Burzenski, Leonard D. Shultz, Michael V. Wiles

**Affiliations:** The Jackson Laboratory, Bar Harbor, ME 04609 USA

**Keywords:** RHBDF2, amphiregulin, metalloproteases, ADAM17, ectodomain shedding, epidermal growth factor receptor, proliferative skin disease, epithelial cancer

## Abstract

The epidermal growth factor (EGF)-receptor ligand amphiregulin (AREG) is a potent growth factor implicated in proliferative skin diseases and in primary and metastatic epithelial cancers. AREG *in vitro*, synthesized as a pro-peptide, requires conversion to an active peptide by metalloproteases by a process known as ectodomain shedding. Although, (Adam17) a disintegrin and metalloprotease 17 is implicated in ectodomain shedding of AREG, it remains to be established *in vivo* whether ADAM17 contributes to AREG shedding. In the present study, using a curly bare (*Rhbdf2^cub^*) mouse model that shows loss-of-hair, enlarged sebaceous glands, and rapid cutaneous wound-healing phenotypes mediated by enhanced *Areg* mRNA and protein levels, we sought to identify the principal ectodomain sheddase of AREG. To this end, we generated *Rhbdf2^cub^* mice lacking ADAM17 specifically in the skin and examined the above phenotypes of *Rhbdf2^cub^* mice. We find that ADAM17 deficiency in the skin of *Rhbdf2^cub^* mice restores a full hair coat, prevents sebaceous-gland enlargement, and impairs the rapid wound-healing phenotype observed in *Rhbdf2^cub^* mice. Furthermore, *in vitro*, stimulated shedding of AREG is abolished in *Rhbdf2^cub^* mouse embryonic keratinocytes lacking ADAM17. Thus, our data demonstrate that ADAM17 is the major ectodomain sheddase of AREG.

## Introduction

The epidermal growth factor receptor (EGFR) pathway plays a major role in normal development, and in multiple diseases including epithelial cancers and chronic obstructive pulmonary disease, and in liver diseases [1-5]. A critical step in regulating this pathway is ectodomain shedding of type-1 transmembrane EGFR ligands from the cell surface by membrane-anchored metalloproteases [6]. For instance, type-1 transmembrane EGFR ligands, including amphiregulin (AREG), transforming growth factor alpha (TGFα), epidermal growth factor (EGF), and heparin-binding EGF (HB-EGF) are produced as inactive pro-peptides. In the ectodomain shedding process, ADAMs (a disintegrin and metalloproteases) cleave pro-peptides to release soluble peptides, leading to activation of the EGFR signaling pathway [7, 8].

Among the multiple ADAMs studied (ADAM8, −9, −10, −12, −15, −17, and −19), ADAM10 and ADAM17 have emerged as key sheddases of the EGFR ligands EGF, betacellulin, HB-EGF, and TGFA [9, 10]. In culture, ectodomain shedding assays using mouse embryonic fibroblasts (mEFs) lacking ADAM10 or ADAM17 show impaired shedding of EGF and betacellulin, and HB-EGF or TGFA, respectively [11]. In line with these findings, ADAM17 knockout mice show defects in cardiac valve and eyelid development [8, 12], defects that are also observed in mice deficient in HB-EGF ( *Hbegf^-/-^* mice) and in TGFA (*Tgfa^-/-^* mice), respectively [12-14]. Furthermore, using loss-of-function experiments in mEFs, Sahin et al. demonstrated that both constitutive and stimulated ectodomain shedding of EGFR ligands, including EGF, epiregulin, betacellulin, HB-EGF, TGFA, and AREG, are unaltered in the absence of ADAM8, −9, −12, −15, and, −19 [9]. Thus, substantial literature suggests that ADAM10 and ADAM17 have essential, but distinct, roles in shedding of EGFR ligands. However, the specific metalloproteases contributing to the ectodomain shedding of AREG, a potent growth factor implicated in proliferative skin diseases, and primary and metastatic epithelial cancers [15-17], remain to be determined *in vivo*. Although results of *in vitro* studies have suggested ADAM17 as a key sheddase, ADAM8-, ADAM15-, and Batimastat (broad metalloprotease inhibitor)-sensitive metalloproteases have also been implicated in AREG shedding *in vitro* [11]. Understanding of the sheddase mechanisms for AREG is critical for development of more effective therapies for diseases associated with this growth factor.

To determine whether ADAM17 is the key sheddase of AREG, we utilized the curly bare (*Rhbdf2^cub^*) gain-of-function mouse mutation. Homozygosity for this spontaneous mutation in the *Rhbdf2* gene augments *Areg* mRNA and protein levels and results in alopecia, sebaceous-gland enlargement, and rapid-wound-healing phenotypes through enhanced secretion of AREG and subsequent hyperactivation of the EGFR pathway [18]. Furthermore, AREG deficiency in *Rhbdf2^cub/cub^* mice prevents the alopecia, sebaceous gland enlargement, and rapid wound-healing phenotypes, suggesting that AREG is the primary mediator of the *Rhbdf2^cub^* phenotype [18]. Thus, the *Rhbdf2^cub^* mouse mutation provides a powerful *in vivo* model system that allows us to examine the physiological role of ADAM17 in ectodomain shedding of AREG and in AREG-mediated downstream events, including wound healing.

Here, we demonstrate that conditional deletion of ADAM17 in the skin of *Rhbdf2^cub/cub^* mice impairs the AREG-mediated hair, sebaceous gland, and wound-healing phenotypes observed in these mice. We also demonstrate that ADAM17 deficiency significantly abolishes both stimulated and unstimulated shedding of AREG in *Rhbdf2^cub/cub^* mouse embryonic keratinocytes (MEKs). Lastly, we show that enhanced shedding of AREG suppresses ADAM17 activity, suggesting that ADAM17 is a key ectodomain sheddase of AREG.

## Materials and Methods

### Animals

All animal work conformed to regulations in the Guide for the Care And Use of Laboratory Animals (Institute of Laboratory Animal Resources, National Research Council, National Academy of Sciences, 8th edition, 2011). Euthanasia was performed in a manner consistent with the 2013 recommendations of the American Veterinary Medical Association (AVMA) *Guidelines on Euthanasia*. All individuals working with animals in this project read and adhered to The Jackson Laboratory policy, POL.AWC.025 *Euthanasia in Animal Experiments Involving Pain, Distress, or Illness.* The *Rhbdf2^cub/cub^, Rhbdf2^-/-^*, and *Rhbdf2^cub/cub^ Areg^-/-^* mice are maintained on the C57BL/6J genetic background, and *Adam17^flox/flox^* and *Adam17^flox/flox^ K14-Cre* mice are of mixed genetic background [19]. *We* generated *Rhbdf2^cub/cub^ *ADAM17^flox/flox^* K14-Cre* mice by crossing female *Rhbdf2^+/cub^*, *Adam17^+/flox^*, *K14-Cre* mice with male *Rhbdf2^cub/cub^*, *Adam17^flox/flox^* mice. *Areg^Mcub/Mcub^* mice are referred to as *Areg^-/-^* mice in this manuscript [18]. Mice were maintained under modified barrier conditions on a 12h light and 12h dark cycle with constant temperature and humidity. The Animal Care and Use Committee at The Jackson Laboratory approved all of the experimental procedures.

### Histology

Mice were euthanized by CO_2_ asphyxiation followed by open chest necropsy, a secondary method of euthanasia. Dorsal skin was removed, fixed in 10% neutral buffered formalin for 24 hours, processed routinely, embedded in paraffin, sectioned and stained, with hematoxylin and eosin (H&E).

### Isolation of primary keratinocytes

For isolation of mouse embryonic keratinocytes (MEKs), skin from embryonic day 18 mouse embryos was incubated overnight in Neutral Protease at 4°C. Following separation of the epidermis from the dermis, the epidermis was placed in Petri dishes containing trypsin (# 12563029,ThermoFisher Scientific) and allowed to incubate for 30 min at room temperature. After blocking trypsin activity with soybean trypsin inhibitor (# R007100, ThermoFisher Scientific), cells were grown in KBM-2 medium (# CC-3107, Lonza) supplemented with antibiotic/antimycotic.

### Measurement of amphiregulin protein levels

AREG levels in the cell culture supernatant were measured via ELISA as described previously [18]. Briefly, 100 µL of cell-culture supernatant was added to capture antibody-pre-coated plates and incubated for 2 h at room temperature (RT). After three washes, 100 µL of the detection antibody was added to each well and incubated for an additional 2 h at RT. Following three washes, 100 µL of Streptavidin-HRP was added to each well and incubated at RT for 20 min, before adding 100 µL of substrate solution (20 min incubation) and 50 µL of stop solution. A spectrophotometer (SpectraMax 190, Molecular Devices) was used to determine the optical density.

### Dextran sulfate sodium (DSS)-induced colitis

*Rhbdf2^+/+^*, *Rhbdf2^cub/cub^*, and *Rhbdf2^cub/cub^ Areg^-/-^* mice were exposed to 2% DSS (MP Biomedicals) for 7 days via drinking water and their body weights were recorded each day for 10 days. The percentage of change in body weight for each mouse strain was documented daily and the differences between the strains were analyzed. After 10 days, mice were euthanized and their colons were examined for hallmarks of colitis.

### SDS/PAGE and immunoblotting

These assays were performed as described previously (10). Briefly, MEFs grown in 6-well dishes were lysed with RIPA buffer (Cell Signaling, MA), and protein concentrations were determined using a Qubit Fluorometer (Life Technologies). After loading 35 µg of protein onto 4-20% (wt/vol) precast gels (Lonza), proteins were transferred to a PVDF membrane followed by blocking with 5% milk for 1 hr at RT. Membranes were then exposed to ADAM17-specific antibody (1:1000, ab2051, Abcam, MA) for 2 hr at RT. Subsequently, membranes were washed in TBST for 2 hr, exposed to secondary antibody (1:5,000, Santa Cruz Biotechnology) for 1 hr at RT, and washed for 2 hr in TBST. Membranes were then exposed to Luminol reagent (Santa Cruz Biotechnology) for 3 min and visualized using a gel doc system (G:BOX F3, Syngene).

### Statistical analysis

One-way ANOVA was used for comparison of several groups using Prism v7 software (GraphPad). A p-value < 0.05 was considered statistically significant. Data represent mean ± SD.

## Results

### The hair-loss, sebaceous-gland enlargement, and rapid wound-healing phenotypes of *Rhbdf2^cub^* mice are mediated through ADAM17

To determine whether ADAM17 is essential for the hair-loss and enlarged sebaceous gland phenotypes exhibited by *Rhbdf2^cub/cub^* mice, we generated *Rhbdf2^cub/cub^* mice lacking ADAM17 in skin, by crossing *Rhbdf2^cub/cub^* mice with *Adam17^flox/flox^ K14-Cre* mice, and studied the phenotypes of second-generation offspring. We noted that ADAM17 acts as a genetic modifier of *Rhbdf2^cub/cub^* mice—*Rhbdf2^cub/cub^ Adam17^flox/flox^ K14-Cre* mice display a full hair coat, in contrast to the complete hair loss in *Rhbdf2^cub/cub^* mice (Fig. 1A). We next performed histopathological examination of truncal skin from *Rhbdf2^cub/cub^* (Fig. 1B, a and b) and *Rhbdf2^cub/cub^ *ADAM17^flox/flox^* K14-Cre* (Fig. 1B, c and d) mice at 3 weeks of age. Although, gross examination of *Rhbdf2^cub/cub^ *ADAM17^flox/flox^* K14-Cre* mice showed a full hair coat (Fig.1A, 3), histological examination revealed mild follicular dystrophy (arrowhead) in *Rhbdf2^cub/cub^ *ADAM17^flox/flox^* K14-Cre* mice compared with extreme follicular dystrophy in *Rhbdf2^cub/cub^* mice (arrows). Additionally, mice of both strains exhibited hyperkeratosis (H) and thickened epidermis (E), whereas enlargement of sebaceous glands (*) was observed only in *Rhbdf2^cub/cub^* mice, suggesting that deletion of ADAM17 partially reverses the hair-loss and sebaceous gland phenotypes of *Rhbdf2^cub/cub^* mice.

**Figure 1.**
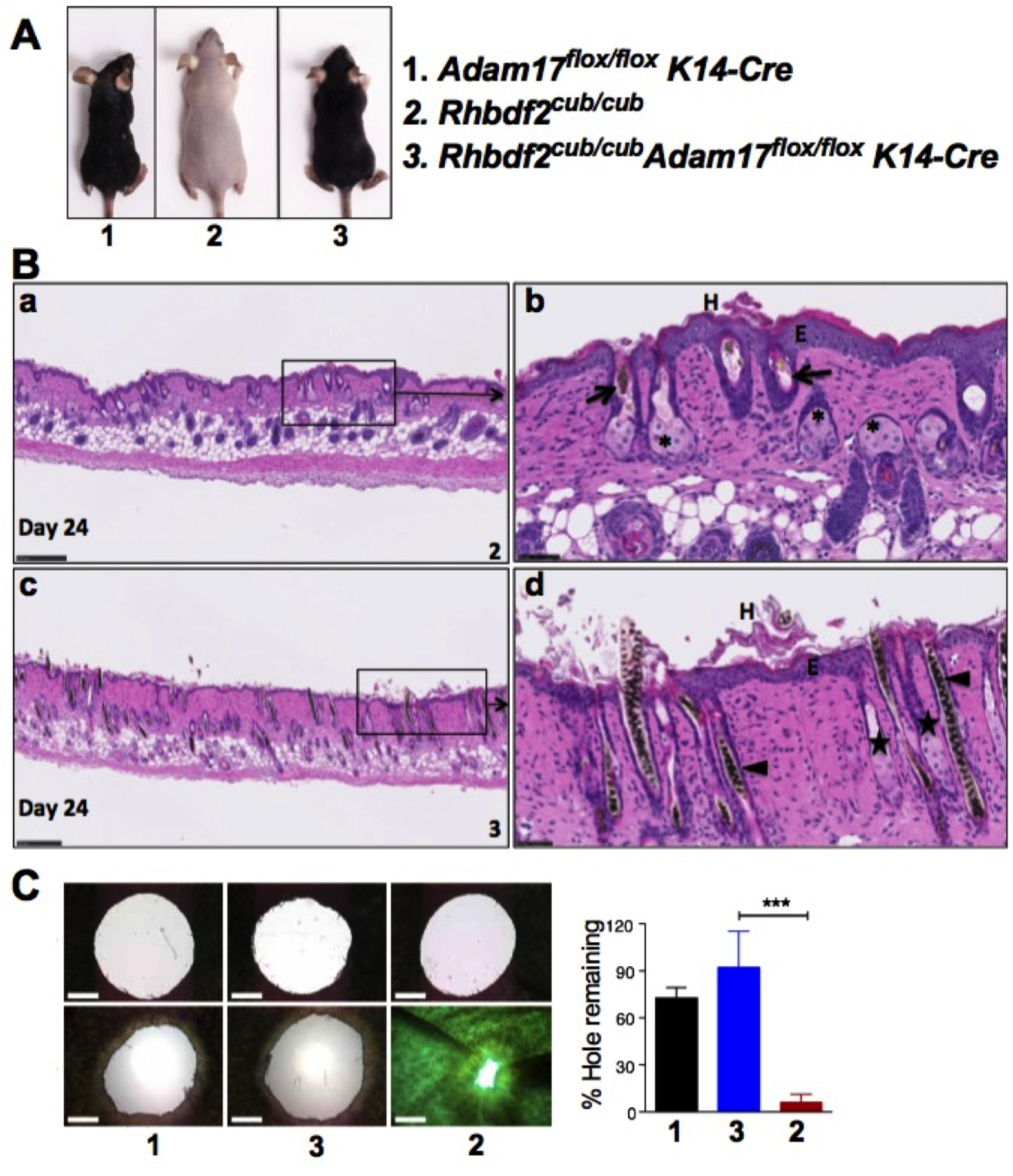
A. Conditional deletion of ADAM17 in the skin of *Rhbdf2^cub/cub^* mice restores hair growth. Also, note the relatively smaller size of the *Rhbdf2^cub/cub^ *ADAM17^flox/flox^* K14-Cre* (3) and *ADAM17^flox/flox^ K14-Cre* (1) mice compared with *Rhbdf2^cub/cub^* (2) mice. B. (a and b) Truncal skin sections of *Rhbdf2^cub/cub^* mice (2) displaying extreme follicular dystrophy (arrow), thickened epidermis (E), enlarged sebaceous glands (*), and hyperkeratosis (H). (c and d) Although *Rhbdf2^cub/cub^ *ADAM17^flox/flox^* K14-Cre* mice exhibit a full hair coat, histological analysis of truncal skin sections of *Rhbdf2^cub/cub^ *ADAM17^flox/flox^* K14-Cre* mice (3) revealed mild follicular dystrophy (arrow heads); however, there was no evidence of sebaceous gland hyperplasia (H). Scale bars: 250 µm (a and c) and 50 µm (b and d). C. Healing of ear tissue in six- to eight-week-old female *ADAM17^flox/flox^ K14-Cre* (1), *Rhbdf2^cub^ *ADAM17^flox/flox^* K14-Cre* (3), and *Rhbdf2^cub/cub^* (2) mice (n=3 per group; representative images are shown) at 0 and 14 days post-wounding. Magnification = 4X; Scale bars = 1mm); Quantification of ear hole closures on day 14; ***p<0.001.

We next examined the rapid wound-healing phenotype of *Rhbdf2^cub/cub^* mice. The *Rhbdf2^cub^* mutation induces a rapid wound-healing phenotype through enhanced secretion of AREG [18, 20]; when we punched 2-mm through-and-through holes in the ear pinnae of *Rhbdf2^cub/cub^* mice, within two weeks ear-hole closure of more than 90% was observed in *Rhbdf2^cub/cub^* mice, in contrast to approximately 20% ear-hole closure in *Rhbdf2^+/+^* mice [18]. Here, we wanted to determine whether the wound-healing phenotype in *Rhbdf2^cub/cub^* mice requires ADAM17. Using the above-mentioned ear-hole closure assay, we tested the wound-healing phenotype of *Rhbdf2^cub/cub^ Adam17^flox/flox^ K14-Cre* mice and compared it to those of *Adam17^flox/flox^ K14-Cre* and *Rhbdf2^cub/cub^* mice. Not surprisingly, impairment of wound healing was similar in *Adam17^flox/flox^ K14-Cre* mice (Fig. 1C, left column) and *Rhbdf2^cub/cub^ Adam17^flox/flox^ K14-Cre* mice (Fig. 1C, middle column), whereas *Rhbdf2^cub/cub^* mice showed the rapid wound-healing phenotype (Fig. 1C, right column). Collectively, these data suggest that loss of ADAM17 in the skin of *Rhbdf2^cub/cub^* mice modifies the hair-loss phenotype and restores a full hair coat, and diminishes the wound-healing phenotype.

### Loss of ADAM17 specifically in the skin causes dermatitis and myeloproliferative disease in *Rhbdf2^cub/cub^* mice

A previous study showed that sustained deficiency of ADAM17 in the epidermis of wildtype mice results in epidermal barrier defects, and subsequently dermatitis and myeloproliferative disease; i.e., a significant increase in the myeloid-cell infiltration [19]. Thus, to determine whether ADAM17 deficiency also causes dermatitis in *Rhbdf2^cub/cub^* mice, we examined the skin phenotype of *Rhbdf2^cub/cub^ *ADAM17^flox/flox^* K14-Cre* mice in comparison with that of *ADAM17^flox/flox^ K14-Cre* and *ADAM17^flox/flox^* control mice. The skin of *Rhbdf2^cub/cub^ ADAM17^flox/flox^ K14-Cre* mice displayed noticeable scaling (Fig. 2A, left), indistinguishable from the phenotype of *ADAM17^flox/flox^ K14-Cre* mice (Fig. 2A, right). Furthermore, histological examination of H&E sections revealed a thicker hypodermis (H) and a thinner dermis (D) in *ADAM17^flox/flox^* control mice (Fig.2B.a) in contrast to a thinner hypodermis and a thicker dermis in both *ADAM17^flox/flox^ K14-Cre* (Fig.2B.b) *and Rhbdf2^cub/cub^ ADAM17^flox/flox^ K14-Cre* (Fig.2B.c) mice. Additionally, we observed epidermal thickening (asterisk), hyperkeratosis (arrow head), and considerable infiltration of inflammatory cells, including macrophages and neutrophils (arrows), in the dermis of *ADAM17^flox/flox^ K14-Cre* mice (Fig. 2B.e) and *Rhbdf2^cub/cub^ *ADAM17^flox/flox^* K14-Cre* mice (Fig. 2B.f), in contrast to *ADAM17^flox/flox^* control mice (Fig.2B.d), which did not manifest any indication of skin disease.

**Figure 2.**
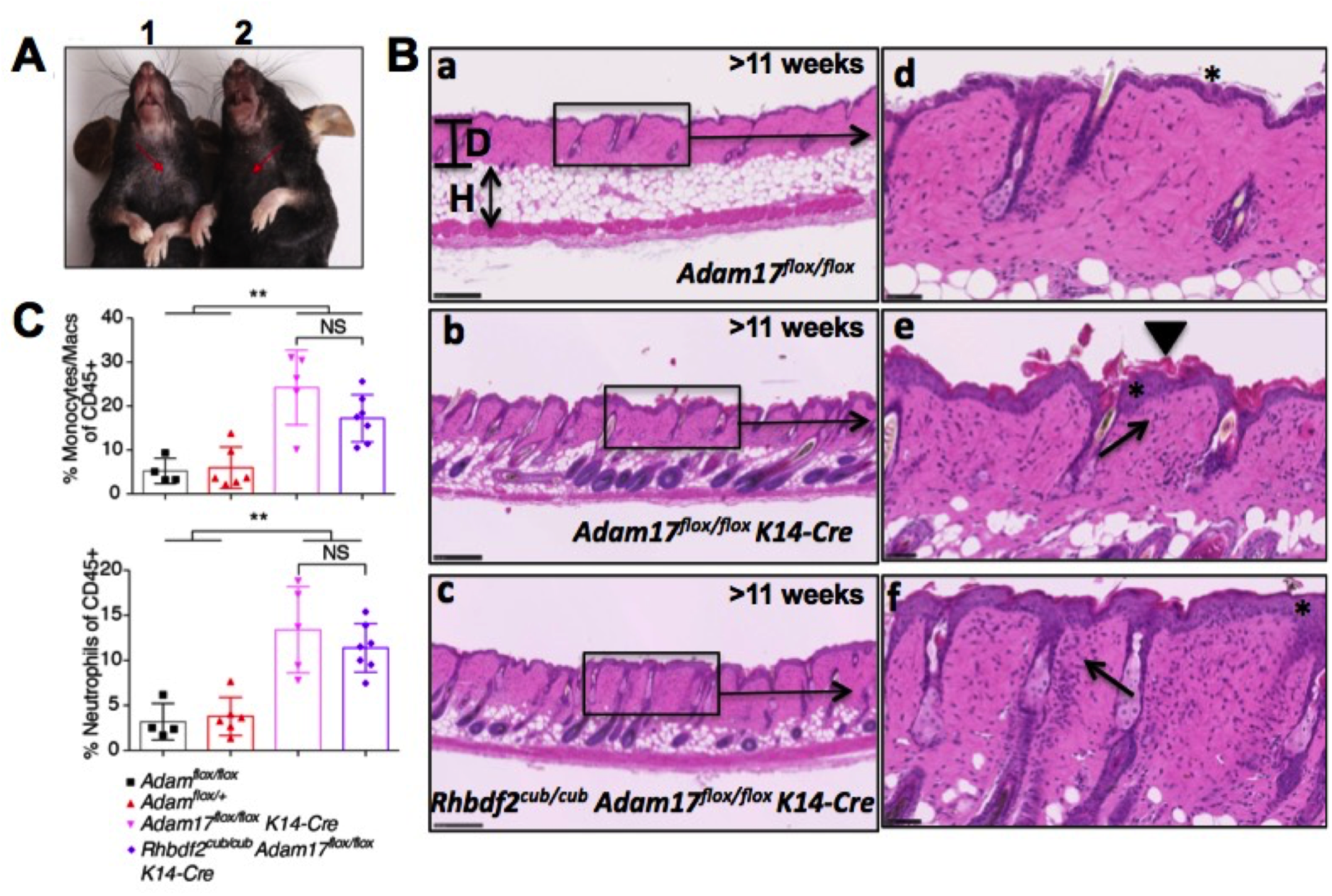
The role of ADAM17 in regulating the skin barrier has been established previously (22)—deletion of ADAM17 in the skin results in epidermal defects and dermatitis. A similar phenotype—dry scaly skin (arrows)—was observed in both *Adam17^flox/flox^ K14-Cre* (1) and *Rhbdf2^cub/cub^ *ADAM17^flox/flox^* K14-Cre* (2) mice. B. Deletion of ADAM17 in the skin results in epidermal defects and dermatitis. A similar dermatitis-like phenotype, including epidermal thickening (asterisk), hyperkeratosis (arrow head), and considerable infiltration of inflammatory cells (arrows) was observed in both *Adam17^flox/flox^ K14-Cre* (b and e) and *Rhbdf2^cub/cub^ *ADAM17^flox/flox^* K14-Cre* (c and f) mice, compared to the normal skin in *Adam17^flox/flox^* control mice (a and d). Note the relatively thicker dermis (D) and thinner hypodermis (H) in both *Adam17^flox/flox^ K14-Cre* (b) and *Rhbdf2^cub/cub^ *ADAM17^flox/flox^* K14-Cre* (c) mice, compared to the thinner dermis and thicker hypodermis in *Adam17^flox/flox^* control mice (a). Scale bars: 250 µm (low magnification) and 50 µm (high magnification). C. Both *ADAM17^flox/flox^ K14-Cre* and *Rhbdf2^cub/cub^ *ADAM17^flox/flox^* K14-Cre* mice develop myeloproliferative disease, evidenced by the increased percentage of macrophages (top) and neutrophils (bottom) in the spleens of these mice compared with control mice (*ADAM17^flox/flox^* and *ADAM17^flox/+^* mice).

Next, using flow cytometry analyses we determined whether there was any indication of myeloproliferative disease in *Rhbdf2^cub/cub^ *ADAM17^flox/flox^* K14-Cre* mice by quantifying the differences in the percentages of splenic macrophages (Fig. 2C, top panel) and neutrophils (Fig. 2C, bottom panel) between *ADAM17^flox/flox^ K14-Cre* and *Rhbdf2^cub/cub^ *ADAM17^flox/flox^* K14-Cre* mice. Compared with control mice (*ADAM17^flox/flox^* and *ADAM17^flox/+^* mice), we observed significantly higher percentages of macrophages and neutrophils in both of *ADAM17^flox/flox^ K14-Cre* and *Rhbdf2^cub/cub^ *ADAM17^flox/flox^* K14-Cre* mice, suggesting that loss of ADAM17 specifically in the skin results in considerable myeloproliferation in *Rhbdf2^cub/cub^* mice. Taken together, our results indicate that lack of ADAM17 in the skin results in dermatitis and myeloproliferative disease, which validates previous findings by Franzke et al. that ADAM17 maintains the skin barrier [19]. Moreover, our results showing development of a similar overt skin phenotype observed by Franzke et al. and restoration of hair growth in *Rhbdf2^cub/cub^* mice lacking ADAM17 implicate ADAM17 as a participant in ectodomain shedding of AREG.

### Ectodomain shedding of AREG requires ADAM17

To determine whether ADAM17 is required for AREG ectodomain shedding, we asked whether loss of ADAM17 in *Rhbdf2^cub/cub^* keratinocytes alters AREG secretion with or without stimulation with 100 nM phorbol-12-myristate-13-acetate (PMA). We also examined AREG-stimulated and –unstimulated secretion in *Rhbdf2^+/+^*, *Rhbdf2^cub/cub^ Areg^-/-^* and *Rhbdf2^-/-^* keratinocytes, for comparison. First, in line with our previous findings [18], we found that both the stimulated (red column) and unstimulated (blue column) secretion of AREG are significantly increased in *Rhbdf2^cub/cub^* keratinocytes compared with control keratinocytes (Fig. 3A; 1, *Rhbdf2^+/+^*; 2,*Rhbdf2^cub/cub^*). Additionally, because *Rhbdf2^cub/cub^ Areg^-/-^* mice do not produce a functional AREG protein [18], as expected, there was no detectable AREG in the culture supernatants of *Rhbdf2^cub/cub^ Areg^-/-^* keratinocytes (Fig. 3A; 3). Furthermore, there was a significant reduction in AREG levels in the culture supernatants of *Rhbdf2^-/-^* keratinocytes with or without stimulation in comparison to those of *Rhbdf2^+/+^* and *Rhbdf2^cub/cub^* keratinocytes (Fig. 3A; 4), suggesting that RHBDF2 is required for both stimulated and unstimulated shedding of AREG.

**Figure 3.**
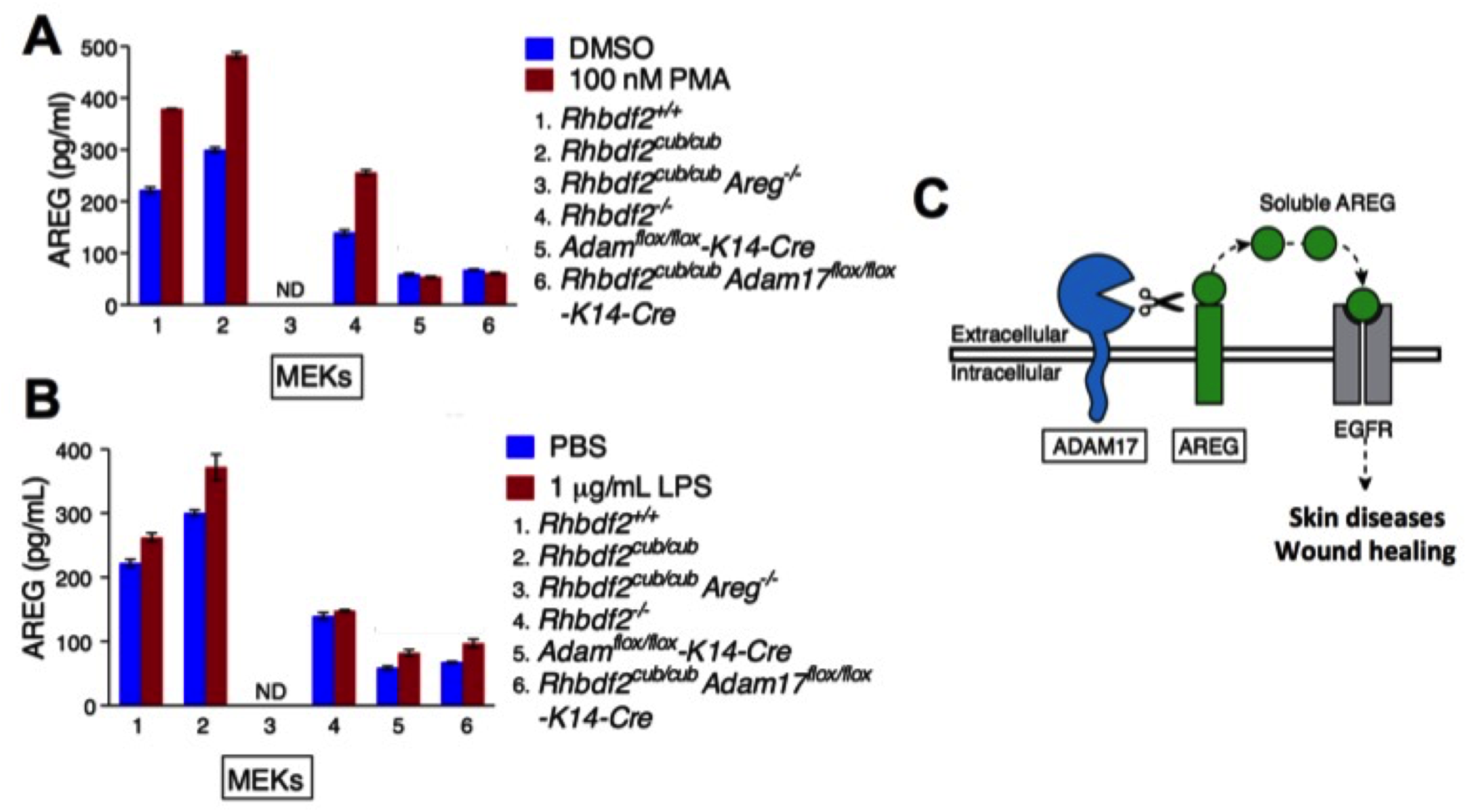
A. ELISA quantitation of cleaved AREG from the supernatant of MEKs after overnight stimulation with either DMSO or 100 nM PMA. ND, not detected; Data represent mean ± SD. B. ELISA quantitation of cleaved AREG from the supernatant of MEKs after overnight stimulation with either PBS or 1 µg/mL LPS. ND, not detected; Data represent mean ± SD. C. Loss of ADAM17 modifies the *Rhbdf2^cub/cub^* hair-loss and ear-punch-closure phenotypes and *in vitro* deficiency of ADAM17 in *Rhbdf2^cub/cub^* keratinocytes prevents stimulated-secretion of AREG, together suggesting that ADAM17 is indispensible for sheddase of AREG.

Second, we observed that, compared to control *Rhbdf2^+/+^* keratinocytes, keratinocytes lacking ADAM17 showed approximately 85% and 75% lower AREG levels in stimulated and unstimulated conditions, respectively (Fig. 3A; 5, *Adam17^flox/flox^ K14-Cre*). Moreover, there was no significant difference in AREG levels between stimulated and unstimulated conditions in *Adam17^flox/flox^ K14-Cre* keratinocytes (Fig. 3A; 5). Third, consistent with the restoration of hair growth and loss of rapid wound-healing phenotypes in *Rhbdf2^cub/cub^ Adam17^flox/flox^ K14-Cre* mice, we observed that *Rhbdf2^cub/cub^ *ADAM17^flox/flox^* K14-Cre* keratinocytes failed to secrete AREG even after stimulation with PMA (Fig. 3A; 6), in contrast to a significant increase in both stimulated and unstimulated *Rhbdf2^cub/cub^* keratinocytes (Fig. 3A; 2), suggesting that the *Rhbdf2^cub^* mutation fails to promote AREG secretion in the absence of ADAM17. Notably, the levels of AREG observed in keratinocytes lacking ADAM17 (Fig. 3A; 5 and 6) can be attributed to constitutive shedding, which does not require ectodomain shedding by metalloproteases [9].

Fourth, similar results were obtained when keratinocytes isolated from the aforementioned strains (1 through 6) of mice were exposed to bacterial endotoxin lipopolysaccharide (LPS) and assayed for AREG levels in the culture supernatants (Fig. 3B). Interestingly, there was a subtle but significant increase in the levels of AREG in *Adam17^flox/flox^ K14-Cre* and *Rhbdf2^cub/cub^ *ADAM17^flox/flox^* K14-Cre* keratinocytes upon stimulation with LPS (Fig. 3B; 5 and 6); however, this could be due to differential regulation of AREG constitutive shedding by PMA versus LPS. Taken together, because loss of ADAM17 significantly abolished stimulated secretion of AREG in *Rhbdf2^cub/cub^* keratinocytes, these data strongly suggest that ADAM17 is essential for ectodomain shedding of AREG (Fig. 3C).

### Enhanced shedding of substrate suppresses ADAM17 sheddase activity

Both our group (Hosur et al., 2014) and Siggs et al. (Siggs et al., 2014) recently observed that ADAM17 activity is reduced in *Rhbdf2^cub/cub^* mice. We sought to validate these findings by assessing the susceptibility of *Rhbdf2^cub/cub^* mice to DSS-induced colitis. Increased susceptibility to DSS-induced colitis is an indicator of reduced ADAM17 activity; conditional ADAM17 knockout mice and ADAM17 hypomorphic mice are highly susceptible to DSS-induced colitis because of impaired EGFR signaling [21, 22]. To examine whether *Rhbdf2^cub/cub^* mice are hypomorphic for ADAM17, in which case they would be more susceptible than control mice to DSS-induced colitis, we provided mice with 2% DSS in drinking water for seven days, followed by water for three days, and recorded the body weight and percentage of change in body weight of *Rhbdf2^+/+^* and *Rhbdf2^cub/cub^* mice. While both *Rhbdf2^+/+^* and *Rhbdf2^cub/cub^* mice experienced weight loss during the seven-day DSS treatment period, *Rhbdf2^cub/cub^* mice continued to experience weight loss during the three-day post-DSS treatment period, while the weight of *Rhbdf2^+/+^* mice stabilized during that period (Fig. 4A). At the end of the ten-day experimental period, *Rhbdf2^cub/cub^* mice had lost significantly more body weight than *Rhbdf2^+/+^* mice (Fig. 4A), suggesting that *Rhbdf2^cub/cub^* mice are more susceptible to DSS-induced colitis. At the conclusion of the ten-day DSS test, experimental animals were euthanized, their colons were removed, and the colonic lengths were compared (Fig. 4B). The colons of *Rhbdf2^+/+^* mice were significantly longer than those of *Rhbdf2^cub/cub^* mice after DSS exposure, further indicating that *Rhbdf2^cub/cub^* mice are more susceptible to DSS-induced colitis. Thus, these data show that ADAM17 activity is significantly reduced in *Rhbdf2^cub/cub^* mice.

**Figure 4.**
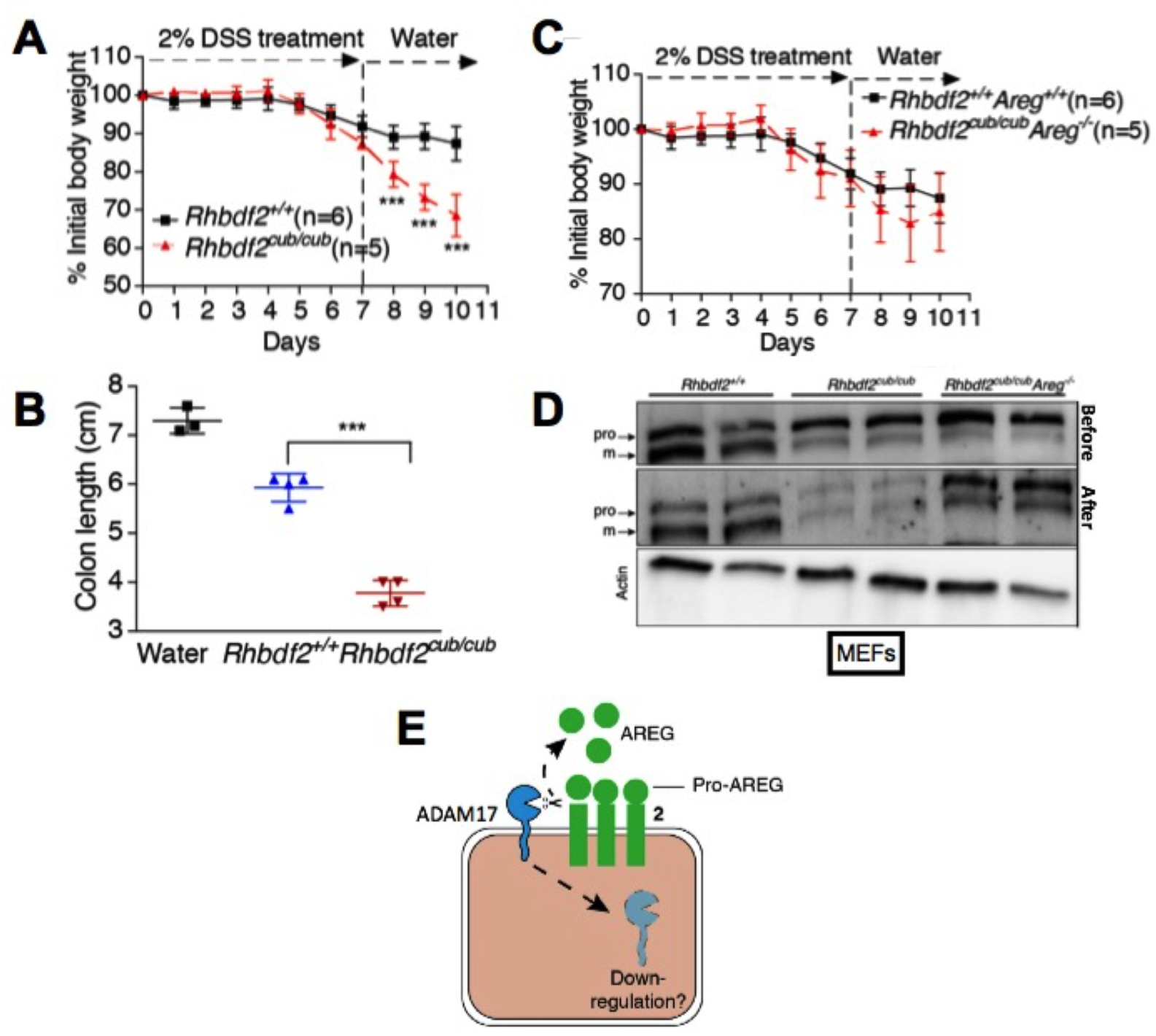
A. Percentage of change in body weights of 8-week-old female *Rhbdf2^+/+^* and *Rhbdf2^cub/cub^* mice over a period of 7 days of exposure to DSS followed by 3 days of water; ***p<0.001. B. Colon lengths from control *Rhbdf2^+/+^* (n=1) and *Rhbdf2^cub/cub^* mice (n=2) on water, and *Rhbdf2^+/+^* (n=4) and *Rhbdf2^cub/cub^* (n=4) mice on DSS, at the end of the experimental period; ***p<0.001. C. Percentage of change in body weights of *Rhbdf2^+/+^ Areg^+/+^* and *Rhbdf2^cub/cub^ Areg^-/-^* mice over a period of 7 days of exposure to DSS followed by 3 days of water. D. Immunoblot analysis of cell lysates obtained from *Rhbdf2^+/+^*, *Rhbdf2^cub/cub^*, and *Rhbdf2^cub/cub^ Areg^-/-^* MEFs before and after stimulation with 100 nM PMA for 24 h. Top row = before stimulation; middle row = after stimulation. Actin served as loading control. pro, pro-ADAM17; m, mature ADAM17. See Supplemental Figure 1 for full blots. E. The observation that ADAM17 activity is significantly reduced in *Rhbdf2^cub/cub^* mice indicates that enhanced shedding of AREG may lead to downregulation or internalization of ADAM17.

To test whether enhanced AREG shedding might reduce ADAM17 levels in *Rhbdf2^cub/cub^* mice, we assessed ADAM17 activity in *Rhbdf2^cub/cub^ Areg^-/-^* mice (Hosur et al., 2014; Johnson et al., 2003) *in vivo*. Here, using the DSS-induced colitis model, we found that whereas *Rhbdf2^cub/cub^* mice are susceptible than *Rhbdf2^+/+^* mice to DSS-induced colitis (Fig. 4A, B), *Rhbdf2^cub/cub^ Areg^-/-^* mice showed no difference in susceptibility to colitis compared with control *Rhbdf2^+/+^ Areg^+/+^* mice (Fig. 4C). The observation that *Rhbdf2^cub/cub^* but not *Rhbdf2^cub/cub^ Areg^-/-^* mice show reduced ADAM17 activity suggests that increased shedding of AREG by ADAM17 in *Rhbdf2^cub/cub^* mice might lead to endocytosis or down-regulation of ADAM17.

To examine stimulated AREG sheddase-induced reduction in ADAM17 protein levels, we performed immunoblot analysis after stimulating MEFs derived from *Rhbdf2^+/+^, Rhbdf2^cub/cub^*, and *Rhbdf2^cub/cub^ Areg^-/-^* mice with PMA for 24h. Whereas ADAM17 levels were lower in unstimulated *Rhbdf2^cub/cub^*, and *Rhbdf2^cub/cub^ Areg^-/-^* MEFs compared with unstimulated *Rhbdf2^+/+^* MEFs, 24h stimulation of MEFs with PMA further significantly reduced ADAM17 levels in *Rhbdf2^cub/cub^* MEFs compared with *Rhbdf2^cub/cub^ Areg^-/-^* MEFs (Fig. 4D), indicating enhanced AREG shedding-induced reduction in ADAM17 levels. Collectively, these data provide strong evidence that *Rhbdf2^cub/cub^* mice are hypomorphic for ADAM17 due to enhanced AREG shedding-induced internalization of ADAM17 (Fig. 4E).

## Discussion

AREG plays an important role in pathological processes, including psoriasis induction [23, 24], cancer progression, and resistance to chemotherapy and anti-EGFR therapies [15, 25]. For example, AREG has been characterized as a multicrine—autocrine, paracrine, and endocrine (systemic)—growth factor in primary and metastatic epithelial cancers [26-28]. AREG induces its own expression to enable self-sufficiency of growth signals acting through EGFR, via an extracellular autocrine loop [29], suggesting that dysregulation of this loop could lead to overexpression of AREG. Additionally, cancer cells overexpressing AREG can induce neoplastic transformation of neighboring cells through paracrine or endocrine activity [30]. Also, more recently, we showed in mice that AREG underlies the hyperproliferative skin disease tylosis, and that loss of AREG restores the normal skin phenotype in a mouse model of human tylosis [31]. Together, these studies highlight the key role of AREG in several pathological processes, and the potential of AREG depletion as a therapeutic approach in multiple diseases. To develop effective therapeutic strategies targeting AREG, it is important to understand how AREG secretion is regulated *in vivo*.

AREG synthesized as pro-AREG is converted to an active form by metalloproteases. Although several ADAMs have been implicated, a study by Sahin et al. [9] showed that *Adam17^-/-^* MEFs exhibit impaired shedding of AREG, indicating that ADAM17 may be a major sheddase. Comparison of the phenotype of *Adam17*^-/-^ mice with that of *Areg*^-/-^ mice is a potential means of providing support for a role of ADAM17 as the major sheddase of AREG. However, literature suggests that in contrast to *Hbegf^-/-^* and *Tgfa^-/-^* mice [13, 14], *Areg^-/-^* mice are viable and do not present with an overt phenotype, except for defects in mammary gland development during puberty and in nursing [32]. Thus, it remains to be determined whether the phenotype of mice with *Areg* depletion resembles any aspects of the *Adam17^-/-^* phenotype. *Adam17^-/-^* mice exhibit perinatal lethality, limiting the ability to examine mammary gland development and nursing competence phenotypes. Furthermore, although at birth *Adam17^-/-^* pups exhibit stunted growth and development [33], including defective mammary branching, suggesting a role for ADAM17 in shedding of AREG, there is a lack of direct evidence. In the present study, using mouse genetics and *in vitro* ectodomain shedding assays, we sought to determine whether loss of ADAM17 abolishes shedding of AREG *in vivo.* We demonstrate that loss of ADAM17 impairs the AREG-mediated loss-of-hair, sebaceous-gland enlargement, and rapid wound-healing phenotypes observed in *Rhbdf2^cub/cub^* mice. Moreover, we find that conditional deletion of ADAM17 in the skin of *Rhbdf2^cub/cub^* mice significantly inhibits stimulated secretion of AREG in keratinocytes. Together, these results suggest that ADAM17 is essential for ectodomain shedding of AREG.

## Author Contributions

V.H., L.D.S., and M.V.W. designed research; V.H., L.M.B., and M.L.F. performed experiments; V.H., L.M.B., and M.L.F. acquired data; V.H., L.D.S., and M.V.W. analyzed data; and V.H., L.D.S., and M.V.W. wrote the paper.

## Conflict of Interest Statement

The authors declare that no conflict of interest exists.

**Supplemental Figure 1.**
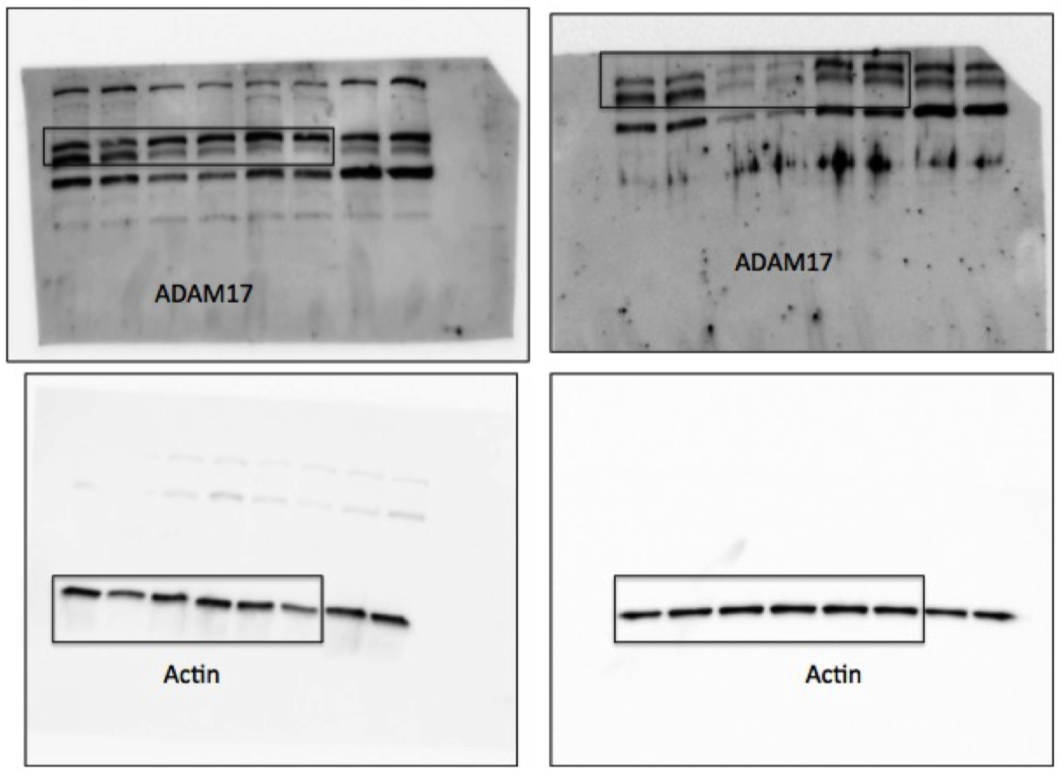

## Acknowledgments

We are grateful to Stephen B. Sampson for critical reading and for providing valuable comments on the manuscript. We also thank Scientific Services at The Jackson Laboratory for assistance with histology (Elaine Bechtel) and flow cytometry (Will Schott and Ted Duffy).

